# High-throughput *in vitro* specificity profiling of natural and high-fidelity CRISPR-Cas9 variants

**DOI:** 10.1101/2020.05.12.091991

**Authors:** Karthik Murugan, Arun S. Seetharam, Andrew J. Severin, Dipali G. Sashital

## Abstract

Cas9 is an RNA-guided endonuclease in the bacterial CRISPR-Cas immune system and a popular tool for genome editing. The most commonly used Cas9 variant, *Streptococcus pyogenes* Cas9 (SpCas9), is relatively non-specific and prone to off-target genome editing. Other Cas9 orthologs and engineered variants of SpCas9 have been reported to be more specific than wild-type (WT) SpCas9. However, systematic comparisons of the cleavage activities of these Cas9 variants have not been reported. In this study, we employed our high-throughput *in vitro* cleavage assay to compare cleavage activities and specificities of two natural Cas9 variants (SpCas9 and *Staphylococcus aureus* Cas9) and three engineered SpCas9 variants (SpCas9 HF1, HypaCas9, and HiFi Cas9). We observed that all Cas9s tested were able to cleave target sequences with up to five mismatches. However, the rate of cleavage of both on-target and off-target sequences varied based on the target sequence and Cas9 variant. For targets with multiple mismatches, SaCas9 and engineered SpCas9 variants are more prone to nicking, while WT SpCas9 creates double-strand breaks (DSB). These differences in cleavage rates and DSB formation may account for the varied specificities observed in genome editing studies. Our analysis reveals mismatch position-dependent, off-target nicking activity of Cas9 variants which have been underreported in previous *in vivo* studies.

## Introduction

Cas9 is the well-studied effector protein of type II CRISPR-Cas (clustered regularly interspaced short palindromic repeats-CRISPR associated) bacterial immune systems (Jiang and Doudna, 2017; Makarova et al., 2020). Cas9 is an endonuclease that uses a dual CRISPR RNA (crRNA) and transactivating-crRNA (tracrRNA) to bind dsDNA targets that are complementary to the guide region of the crRNA and adjacent to a short, conserved protospacer-adjacent motif (PAM) sequence (Gasiunas et al., 2012). Two nuclease domains in Cas9, HNH and RuvC, cut the target and non-target strand respectively, generating a double-stranded break (DSB) in the dsDNA (Jinek et al., 2012) with little post-cleavage trimming (Stephenson et al., 2018). The dual RNAs can be combined into a single guide-RNA (sgRNA) and the targeting region can be varied, making Cas9-sgRNA a readily programmable, two component system for use in various biotechnological applications (Doudna, 2020; Jinek et al., 2012). In particular, DSB formation followed by DNA repair can lead to changes in genomic DNA sequence, enabling genome editing following Cas9 cleavage (Cong et al., 2013; Mali et al., 2013).

Cas9 can tolerate mismatches between the crRNA and the target DNA, which is consistent with its role as a bacterial immune system effector in facilitating defense against rapidly evolving phages (Fu et al., 2016, 2014, 2013; Hsu et al., 2013; Pattanayak et al., 2013). Cas9 generally tolerates multiple mismatches in the PAM-distal region while PAM-proximal “seed” mismatches reduce the cleavage activity. This low fidelity leads to off-target activity when used for genome editing applications, as Cas9 can create DSBs at sites with limited homology to the intended target. Many strategies have been developed to reduce off-target activity of Cas9 (Vakulskas and Behlke, 2019). Among these, various natural and engineered variants of Cas9 have been reported to have reduced off-target cleavage activity. While the commonly used wildtype (WT) *Streptococcus pyogenes* Cas9 (SpCas9) can tolerate multiple mismatches in the target sequence, other naturally occurring Cas9 orthologs from *Staphylococcus aureus* (SaCas9), *Neisseria meningitidis* and *Campylobacter jejuni* are reported to have higher specificity in genome editing compared to SpCas9 (Amrani et al., 2018; Friedland et al., 2015; Kim et al., 2017; Ran et al., 2015). SpCas9 has also been engineered to improve the fidelity of target cleavage activity. Some mutations were designed to reduce DNA target interactions, making the requirement for complete complementary with the crRNA more stringent (Kleinstiver et al., 2016; Slaymaker et al., 2016). Mutations rationally introduced in the REC domain of SpCas9 prevent conformational changes required for nuclease domain activation when a target sequence with mismatches is encountered (Chen et al., 2017; Dagdas et al., 2017). Bacterial screens have also been used to select high-fidelity SpCas9 variants that maintain on-target cleavage but have reduced off-target cleavage activity (Hu et al., 2018; Lee et al., 2018; Vakulskas et al., 2018).

Several methods have been developed to detect and study off-target activities of Cas9 (Tsai et al., 2017, 2015; Vakulskas and Behlke, 2019). However, methods that measure Cas9 off-target editing in eukaryotic cells are limited because cellular factors and DNA accessibility sequester potential cleavage sites (Horlbeck et al., 2016; Yarrington et al., 2018). DNA accessibility can also vary depending on cellular processes which may change the outcome and detection of potential Cas9 off-target editing events. These methods also rely on DSBs in the DNA generated by Cas9 or post-cleavage DNA repair and indel formation which can vary among cell types and experiments (Schmid-Burgk et al., 2020; Tsai et al., 2017, 2015). Differences in the Cas9/sgRNA delivery methods and cell lines have resulted in discrepancies in the reported specificities of high-fidelity Cas9 variants (Chen et al., 2017; Kleinstiver et al., 2016; Vakulskas et al., 2018).

To avoid these pitfalls, specificity studies can be performed *in vitro* to detect the native cleavage activities of Cas9 variants (Fu et al., 2019, 2016, 2014; Huston et al., 2019; Jones et al., 2019; Pattanayak et al., 2013; Zhang et al., 2020). Here, we used a previously established plasmid-based high-throughput *in vitro* cleavage assay to compare the native cleavage specificity of different Cas9 variants (Murugan et al., 2020). Our method enables the detection of target sequences that may be nicked by Cas9. We tested the cleavage activity of two WT Cas9 orthologs, SpCas9 and SaCas9, and three engineered SpCas9 variants, SpCas9 HF1 (Kleinstiver et al., 2016), hyper-accurate Cas9 (HypaCas9) (Chen et al., 2017) and Alt-R ® S.p. HiFi Cas9 (Vakulskas et al., 2018) against two different target library sequences. Each of these three variants represent a version of high-fidelity Cas9 developed via different strategies discussed above. We show that SpCas9 rapidly cleaves target sequences with up to five mismatches. While the high-fidelity Cas9 variants retained cleavage activity against targets with multiple mismatches, they have reduced rates of cleavage compared to SpCas9. High-fidelity Cas9 variants also have higher nicking activity against target sequences with mismatches, resulting in incomplete DSB formation. Overall, our study reveals target-sequence dependent nicking activity that may account for the lower off-target cleavage events observed in genome editing studies with high-fidelity Cas9 variants.

## Results

### Cleavage activity of Cas9 against target library

We sought to compare the cleavage activity and specificity of Cas9 and engineered variants in a high-throughput manner. We performed a previously established *in vitro* plasmid library (pLibrary) cleavage assay with Cas9 and dual RNAs (tracrRNA and crRNA) followed by high-throughput sequencing analysis (Fig. 1A, S1, S2A, B) (Murugan et al., 2020). The pLibraries contained a distribution of target sequences with zero and ten mismatches to the crRNA guide sequence, with a maximum representation of target sequences with two to four mismatches in the library (Fig. S2B). We tested the cleavage activity of WT SpCas9, WT SaCas9 and three high-fidelity variants of SpCas9 – SpCas9 HF1, HypaCas9, and Alt-R ® S.p. HiFi Cas9 (HiFi Cas9) (Chen et al., 2017; Kleinstiver et al., 2016; Vakulskas et al., 2018). The three high-fidelity variants of SpCas9 will be collectively referred to as HF Cas9 hereafter. We used two different crRNA sequences and generated corresponding negatively supercoiled (nSC) plasmids containing the perfect target (pTarget) or target library (pLibrary) (see methods section – Plasmid and nucleic acid preparation) (Fig. S2B). The two crRNA and library sequences were designed based on protospacer 4 sequence from *Streptococcus pyogenes* CRISPR locus (55% G/C) and EMX1 gene target sequence (80% G/C), referred to as pLibrary PS4 and EMX1 respectively.

**Figure 1.**
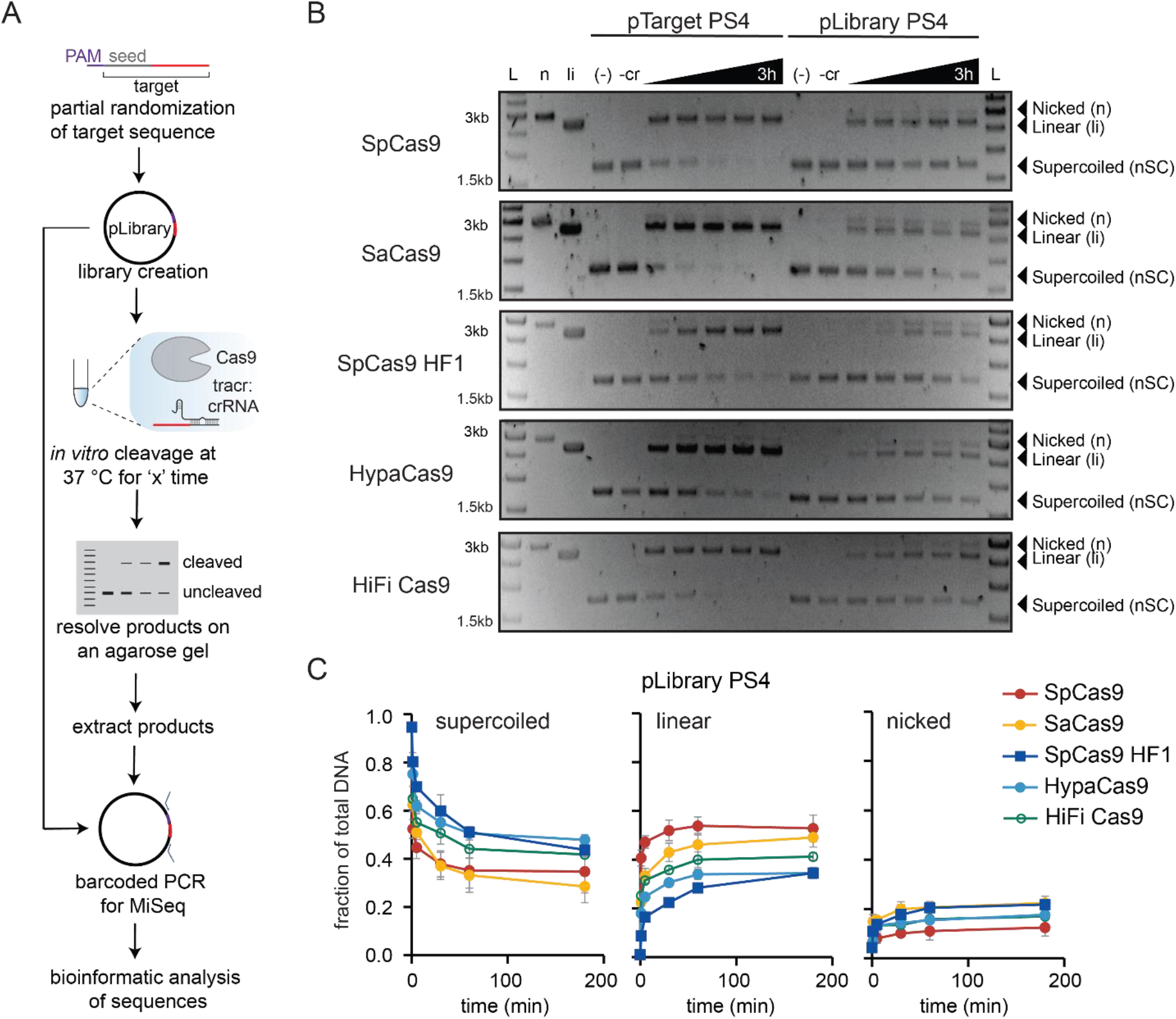
High-throughput *in vitro* analysis of Cas9 mismatch tolerance. (A) Outline and workflow of the high-throughput *in vitro* cleavage assay. (B) Representative agarose gel showing time course cleavage of negatively supercoiled (nSC) plasmid containing a fully matched PS4 target (pTarget PS4, left) and plasmid library PS4 (pLibrary PS4, right) by Cas9 variants, resulting in linear (li) and/or nicked (n) products. Time points at which the samples were collected are 1 min, 5 min, 30 min, 1 h, and 3 h. All controls were performed under the same conditions as the longest time point for the experimental samples. Controls: (−) = pTarget or pLibrary alone incubated at 37 °C for the longest time point in the assay (3 h); (-r) = pTarget or pLibrary incubated with Cas9 only at 37 °C for the longest time point in the assay (3 h); n = Nt.BspQI nicked pUC19; li = BsaI-HF linearized pUC19 (C) Overall cleavage of the pLibrary PS4 by Cas9 indicating the decrease in supercoiled (nSC) pool and appearance of nicked (n) and linear (li) pools over time. The 0 time point is the quantification of the negative control pLibrary (i.e. pLibrary run on a gel after preparation as represented in Fig. S2A). Values plotted represent an average of two replicates. Error bars are SEM.

We used the differential migration of the nicked (n) and linear (li) cleavage products of negatively supercoiled (nSC) dsDNA plasmid on an agarose gel to analyze Cas9 cleavage activity (Oppenheim, 1981). Cleavage rates of pTarget were variable depending both on target sequence and Cas9 variant (Fig. 1B, S2C). SpCas9 HF1 and HypaCas9 cleaved pTarget PS4 relatively slowly (Fig. 1B) but were comparable to SpCas9 and HiFi Cas9 for pTarget EMX1 (Fig. S2C). While cleavage of pTarget PS4 by SaCas9 was comparable to SpCas9, cleavage of pTarget EMX1 by SaCas9 was substantially reduced in comparison to other Cas9 variants (Fig. S2C). Cleavage rates of the pLibrary were consistent with those observed for pTarget. While, SpCas9 HF1 and HypaCas9 cleaved pLibrary PS4 relatively slowly, SaCas9 had the slowest rate of cleavage for pLibrary EMX1 (Fig. 1B, C, S2C, D). SpCas9 rapidly cleaved more than 50% of both negatively supercoiled pLibraries, with the vast majority of product DNA becoming linearized (Fig. 1C, S2D). In contrast, for HF Cas9 variants and SaCas9, we observed accumulation of nicked pool, especially for pLibrary PS4 (Fig. 1C, S2D).

We also checked whether cleavage occurred outside of the target region during pLibrary cleavage by testing the cleavage activity of Cas9 against the empty plasmid backbone without and with the different crRNAs (Fig. S2E). The empty plasmid was minimally cleaved by Cas9-tracrRNA:crRNA, except in the case of SpCas9-EMX1 crRNA where a substantial nicked product was observed at the three hour time point. However, we do not observe similar amounts of nicking of the pLibrary EMX1 by Cas9 (Fig. S2D) and further analysis indicated that pLibrary nicking is, in part, target-sequence dependent (see below).

To determine which sequences were cleaved by Cas9 variants, we extracted the plasmid DNA from the supercoiled and nicked pools, performed barcoded-PCR amplification and multiplexed, high-throughput sequencing (HTS) (Fig. 1A, S1). Target sequences cleaved by Cas9 were depleted from the supercoiled pool while those nicked by Cas9 were enriched in the nicked pool. Although we were unable to sequence the linearized pool using PCR-based HTS, for our analysis, we assumed that target sequences absent in both the supercoiled and nicked pools were linearized. We normalized the counts of the target sequences in the HTS data with the fraction of DNA present in the pool at a given time point (Fig. 1C, S2D), and then to the original library to obtain relative, normalized counts (see methods section – HTS analysis).

We generated mismatch distribution curves to compare the cleavage of target sequences containing varying numbers of mismatches with the crRNA across Cas9 variants at each time point or across time points for each Cas9 variant (Fig. 2, S3). As expected, the perfect target (zero mismatch) was quickly depleted from the supercoiled pool of the pLibrary (Fig. 2A, C, S3A, C). HF Cas9 variants cleaved all target sequences more slowly than SpCas9 for pLibrary PS4 but had comparable cleavage activity for G/C rich pLibrary EMX1. SpCas9 quickly cleaved sequences with up to four mismatches (i.e. target sequences containing four mismatches with the crRNA guide sequence) in the first time point tested for both pLibraries. While HF Cas9 variants cleaved target sequences relatively slowly in pLibrary PS4, they also eventually cleaved target sequences with up to four mismatches (Fig. 2A, C). Sequences with six or more mismatches were not notably depleted from the supercoiled pool indicating that these sequences were not cleaved by Cas9 (Fig. 2A, C, S3A, C).

**Figure 2.**
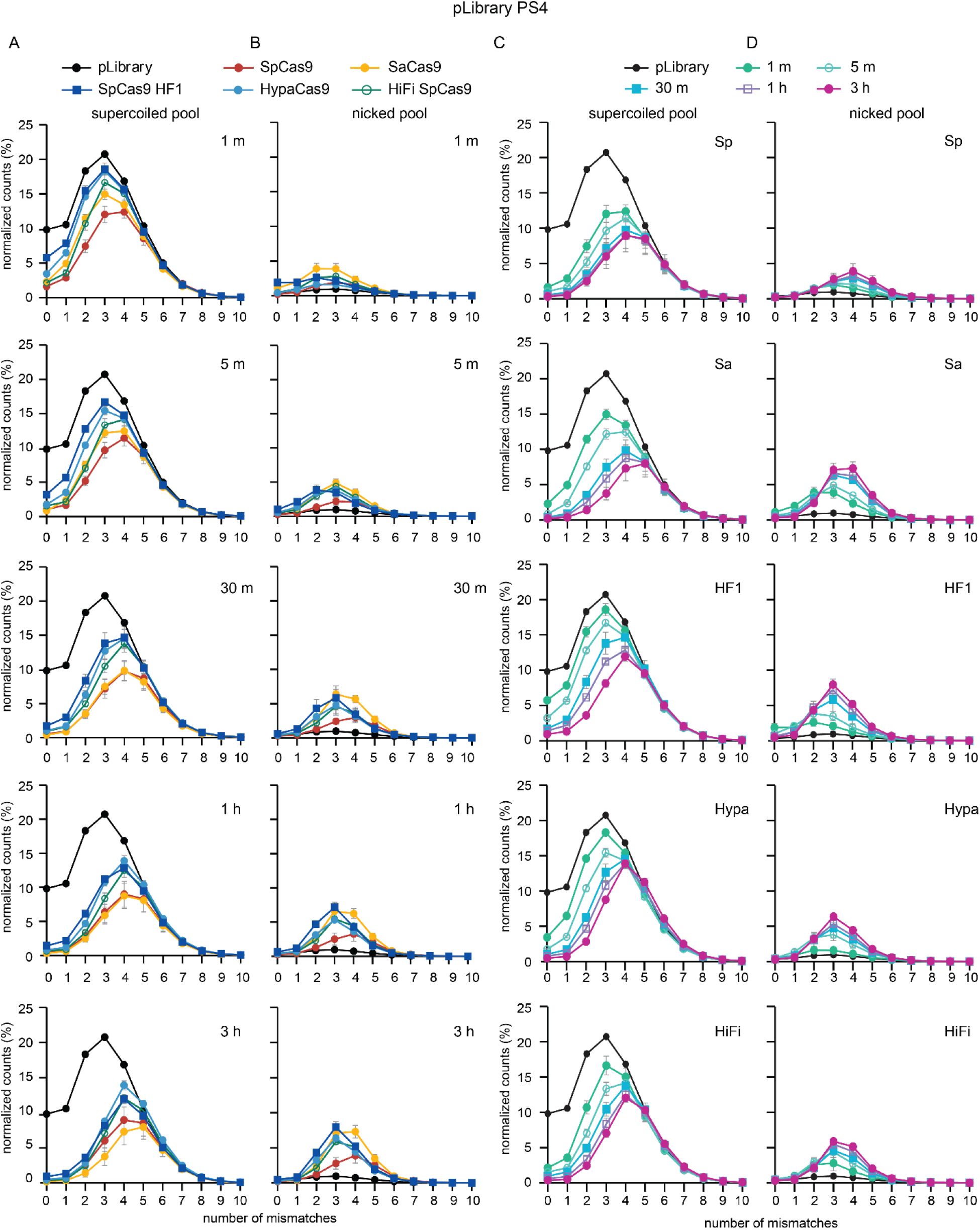
Mismatch distribution curves for Cas9 cleavage activity against pLibrary PS4. Mismatch distribution of (A, C) supercoiled pool and (B, D) nicked pool from pLibrary PS4 when subject to cleavage by different Cas9 variants. The mismatch distribution is plotted for (A,B) each time point or (C, D) each Cas9 variant. Depletion of target sequences from the supercoiled pool indicates cleavage, and enrichment in the nicked pool indicates nicking. The decrease in nicked pool over time indicates linearization of target sequences. Values plotted represent an average of two replicates. Error bars are propagation of SEM. Sp = SpCas9, Sa = SaCas9, HF1 = SpCas9 HF1, Hypa = HypaCas9, HiFi = Alt-R ® S.p. HiFi Cas9.

SaCas9 and HF Cas9 variants nicked target sequences with two to four mismatches, based on the accumulation of these sequences in the nicked pool over time (Fig. 2B, D, S3B, D). Some target sequences with one and two mismatches were initially nicked but subsequently linearized over time. Sequences with three to five mismatches accumulated in the nicked pool for pLibrary PS4 while those in pLibrary EMX1 did not change notably over time (Fig. 2B, D).

### Specificity comparison for Cas9 variants

Our HTS data allows us to compare the overall cleavage efficiency and specificity of the Cas9 variants. We first determined the efficiency of cleavage of the perfect target and targets with multiple mismatches (one to five MM) (Fig. 3A – D) (see methods – HTS analysis). Cleavage efficiencies of the perfect target within pLibrary were similar to those observed for pTarget, with reduced rates for HypaCas9 and SpCas9 HF1 against the PS4 perfect target and reduced rates for SaCas9 against the EMX1 perfect target (Fig. 1B, S2C, 3A, B). Analysis of mismatch targets indicated differences in cleavage efficiencies in comparison to the perfect target (Fig. 3C, D). For example, while HiFi Cas9 cleaved the PS4 perfect target with similar efficiency to SpCas9, we observed a marked reduction in cleavage of PS4 mismatched targets for HiFi Cas9 (Fig. 3C).

**Figure 3.**
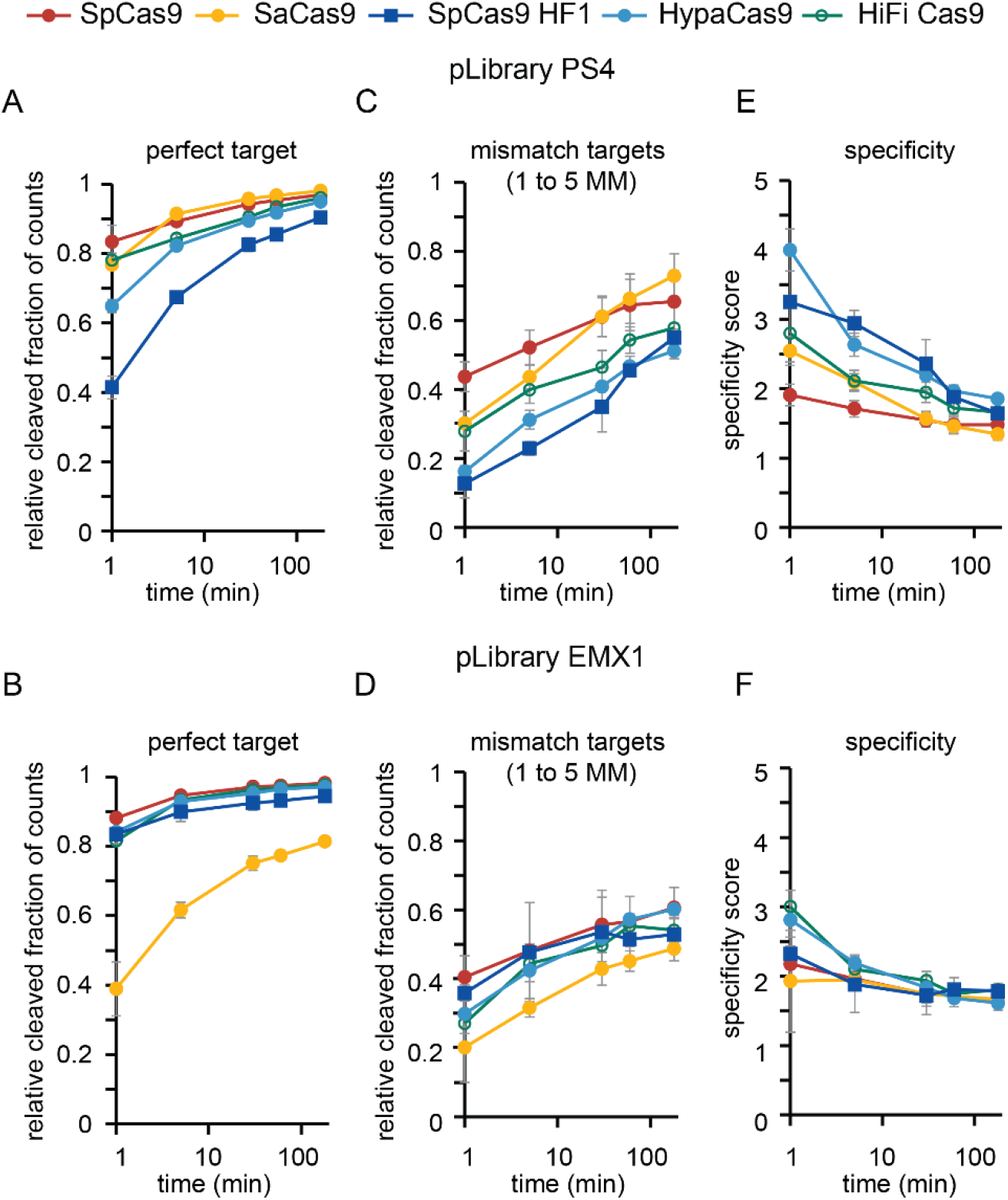
Specificity scores for Cas9 variants. The fractions of the (A, B) perfect target and (C, D) target sequences with 1 to 5 mismatches (MM) that were cleaved by Cas9 variants are plotted versus time for pLibrary (A, C) PS4 and (B, D) EMX1. Specificity scores were calculated for Cas9 cleavage of pLibraries (E) PS4 and (F) EMX1 over the time course of the assay (see methods – HTS analysis). Values plotted represent an average of two replicates. Error bars are propagation of SEM.

To analyze these differences in cleavage efficiencies, we generated a specificity score that reports the relative efficiency of cleavage of on-and off-target sequences over time for the Cas9 variants against the two pLibraries (Fig. 3E, F) (See methods – HTS analysis). Thus, high specificity score values indicate relatively high specificity, while low values indicate relatively low specificity. SpCas9 scored the lowest for both pLibraries, consistent with its expected low specificity. At early time points, specificity scores of Cas9 variants were highly variable. While SaCas9 initially appeared more specific than SpCas9 at early time points for pLibrary PS4, at later time points both orthologs scored similarly. HF Cas9 variants generally scored higher than SpCas9 for pLibrary PS4, with SpCas9 HF1 and HypaCas9 scoring highest (Fig. 3A, B). In contrast, SpCas9 HF1 scored similarly to SpCas9 and SaCas9 for pLibrary EMX1, indicating that increased specificity for this variant may be target dependent. In general, we observed more spread in specificity scores for pLibrary PS4 than for pLibrary EMX1, suggesting that target dependence is universal to specificity of all Cas9 variants. Interestingly, specificity scores for HF Cas9s changed over time and eventually converged with SpCas9 and SaCas9. This change in specificity score over time indicates that prolonged exposure of HF Cas9 variants can lead to off-target cleavage activity.

### Sequence determinants of Cas9 cleavage activity

To characterize the effects of the mismatch position and type on Cas9 cleavage, we analyzed the sequences present in both the supercoiled and nicked pools (Fig. S1). We generated heatmaps showing the relative abundance of target sequences containing one to six mismatches over time (Fig. 4, 5, S4, S5). These heatmaps show the effect of all possible nucleotides at each position of the sequence in the supercoiled pool when present in a target sequence as a single mismatch or the cumulative effect of multiple mismatches (see methods section – HTS analysis).

**Figure 4.**
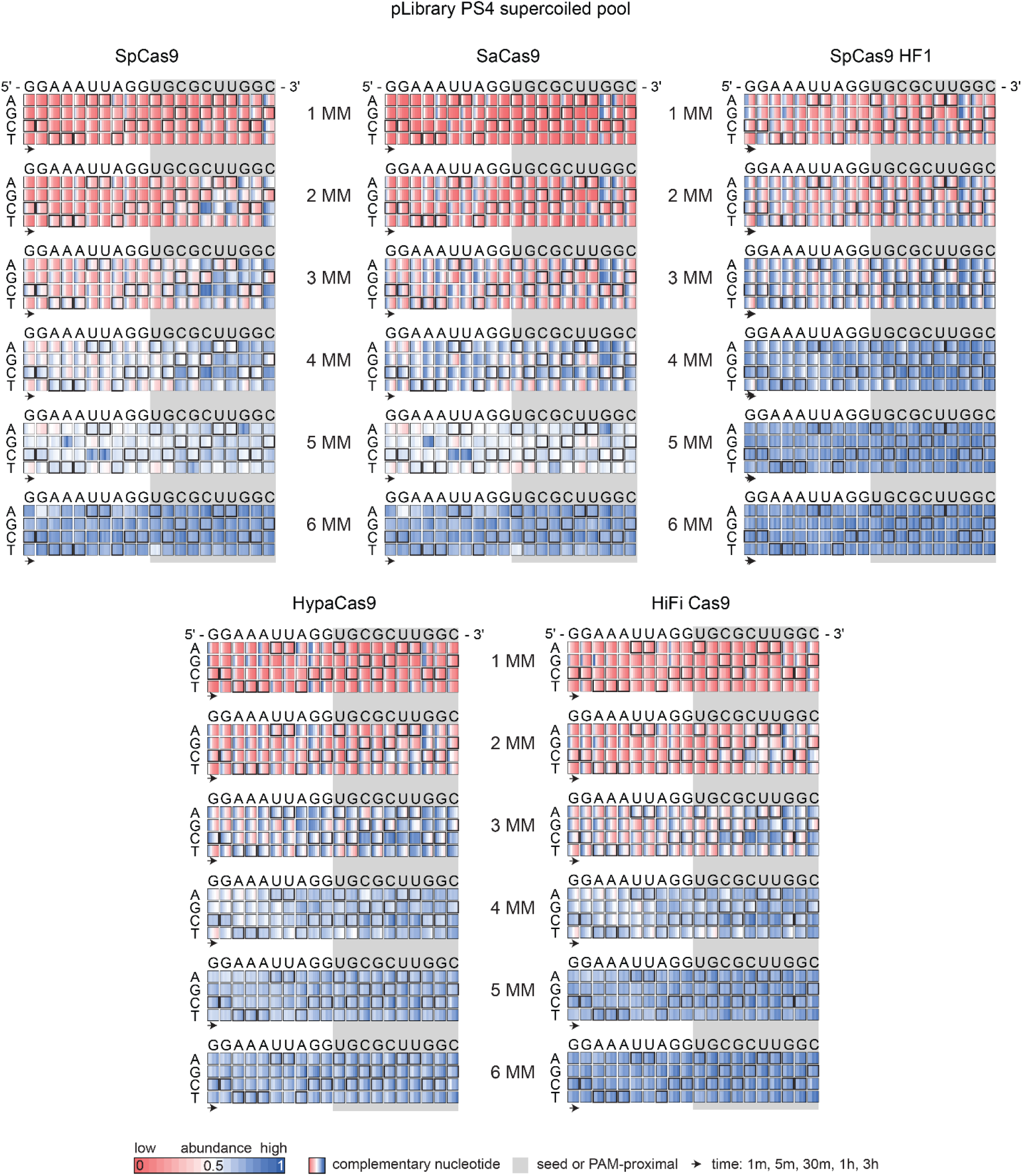
Sequence determinants of Cas9 cleavage activity for pLibrary PS4. Heatmaps showing the relative abundance of different mismatched sequences over time for the supercoiled pool in pLibrary PS4 upon cleavage by Cas9 variants. The targeting region of the crRNA sequence is indicated on the top. The nucleotides on the left side of the heatmaps indicate the potential base pair or mismatches formed. The crRNA-complementary nucleotides are highlighted by bold black boxes in the heatmap which result in Watson–Crick base pairs. The PAM-proximal “seed” sequence is highlighted by the grey box. Each box represents normalized proportion of sequences containing each nucleotide at a given position across time. Values plotted represent an average of two replicates. MM = mismatch.

**Figure 5.**
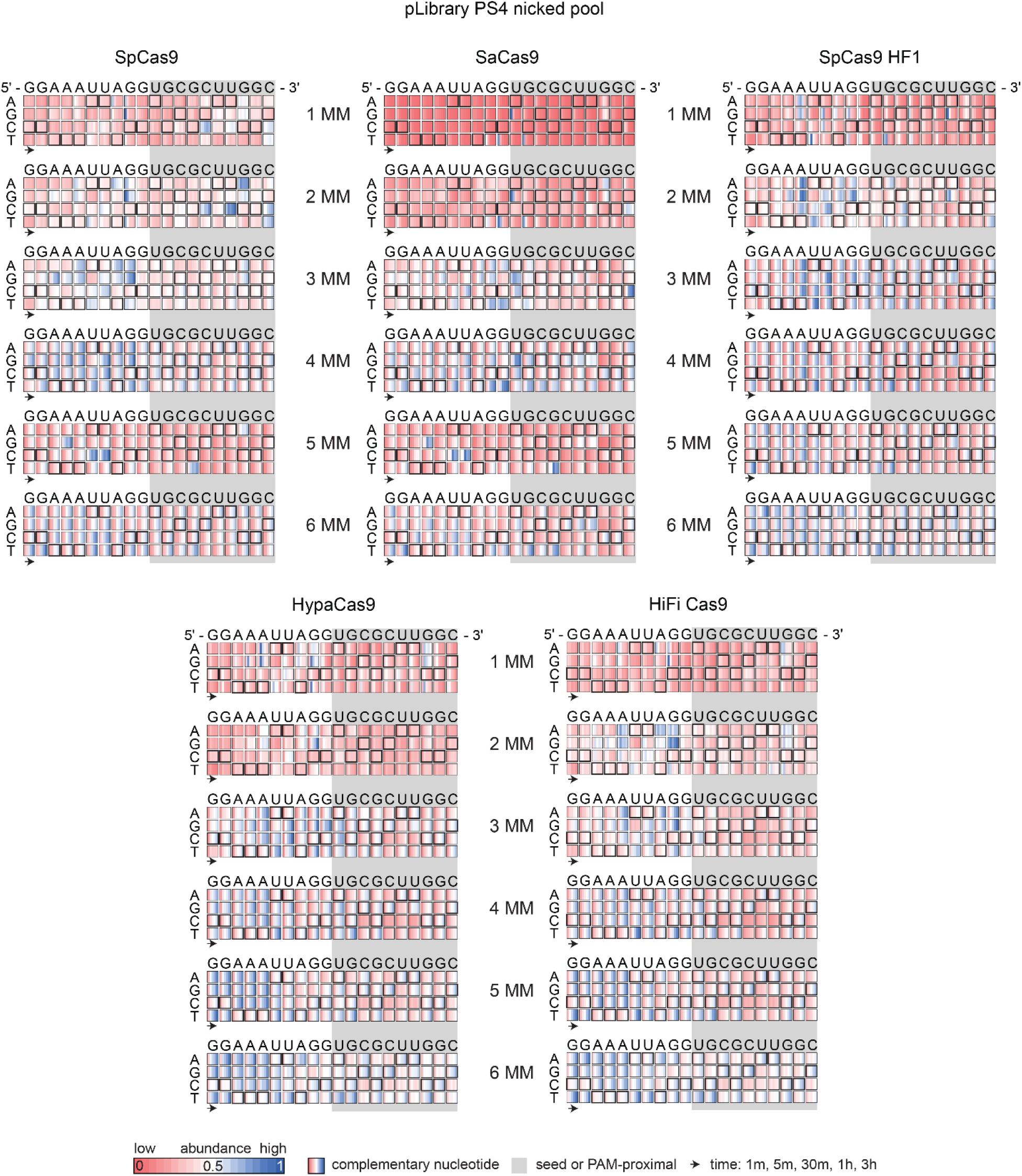
Sequence determinants of Cas9 nicking activity for pLibrary PS4. Heatmaps showing the relative abundance of different mismatched sequences over time for the nicked pool in pLibrary PS4 upon cleavage by Cas9 variants. The targeting region of the crRNA sequence is indicated on the top. The nucleotides on the left side of the heatmaps indicate the potential base pair or mismatches formed. The crRNA-complementary nucleotides are highlighted by bold black boxes in the heatmap which result in Watson–Crick base pairs. The PAM-proximal “seed” sequence is highlighted by the grey box. Each box represents normalized proportion of sequences containing each nucleotide at a given position across time. Values plotted represent an average of two replicates. MM = mismatch.

The heatmaps revealed a strong PAM-proximal “seed” dependent cleavage activity, as expected for all Cas9s, especially for target sequences with two to four mismatches. The seed region is eight to ten nucleotides from the PAM for Cas9 as previously reported (Hsu et al., 2013; Liu et al., 2016; Sternberg et al., 2014) (Fig. 4, 5, S4, S5). Although target sequences with a single mismatch were depleted from the supercoiled pool by SpCas9 for both pLibraries, mismatches in the seed region slowed down the rate of depletion (Fig. 4, S4). For SaCas9 and HF Cas9 variants, single mismatches in the PAM-distal region were generally more deleterious than for SpCas9. A single mismatch throughout the target sequence in pLibrary PS4 was more deleterious for cleavage by SpCas9 HF1 and HypaCas9 (Fig. 4). HF Cas9 variants also better tolerated single mismatches outside the seed in G/C rich pLibrary EMX1 (Fig. S4).

Similarly, SpCas9 was more tolerant of two to four mismatches outside the seed while SaCas9 and HF Cas9 variants were slower at depleting these target sequences from the supercoiled pool (Fig. 4, S4). In pLibrary EMX1, C and G mismatches slow the rate of depletion of these target sequences for all Cas9s (Fig. S4). The heatmaps also revealed the depletion of some target sequences containing five and some with six mismatches in the PAM-distal end, particularly by SpCas9 and SaCas9 in pLibrary PS4 (Fig. 4).

Cas9 also had high target-sequence dependent nicking activity for target sequences with multiple mismatches (Fig. 5, S5). Cas9 nicked and slowly linearized target sequences with mismatches across the length of the target, although we observed a marked effect of PAM-distal mismatches (Fig. 5, S5). In particular, target sequences with two to four mismatches in the PAM-distal region accumulated over time in the nicked pool. The nicked pool of pLibrary PS4 also contained target sequences with five and six mismatches in the PAM-distal region (Fig. 5). These results indicate that SpCas9 is more promiscuous than previously thought where it can also nick target sequences with three to six mismatches that were not recorded in previous *in vitro* and *in vivo* specificity studies (Fu et al., 2019, 2016, 2014, 2013; Hsu et al., 2013; Pattanayak et al., 2013). SaCas9 and HF Cas9 variants are also more prone to nicking sequences with several mismatches in the PAM-distal region. For pLibrary EMX1, HiFi Cas9 eventually linearized some of the target sequences with multiple mismatches (Fig. S5).

### Effect of two mismatches as function of position and separation on Cas9 cleavage activity

Our library was designed to have a high representation of targets sequences containing two mismatches with the crRNA. To further understand the effects of two mismatches on Cas9 cleavage, we generated heatmaps of the relative abundance of these target sequences as a function of mismatch location and distance between the two mismatches (see methods section – HTS analysis).

There are 190 possible combinations in which two mismatches can occur in a 20-nucleotide sequence, in which the two mismatches can be separated by between 0 and 18 nucleotides. We observed that target sequences with two mismatches that are separated by four nucleotides or less are generally depleted slowly from the supercoiled pool by all Cas9 variants (Fig. 6). However, SpCas9 is more tolerant of two mismatches compared to the other Cas9 variants tested, even when the mismatches are spaced close together (Fig. 6A, B). SpCas9 HF1 initially nicked most sequences with two mismatches separated by any distance in both pLibraries. Over time, most target sequences with two mismatches separated by a distance of six or fewer nucleotides remained nicked while sequences with mismatches that were further apart were linearized by SpCas9 HF1 in pLibrary EMX1.

**Figure 6.**
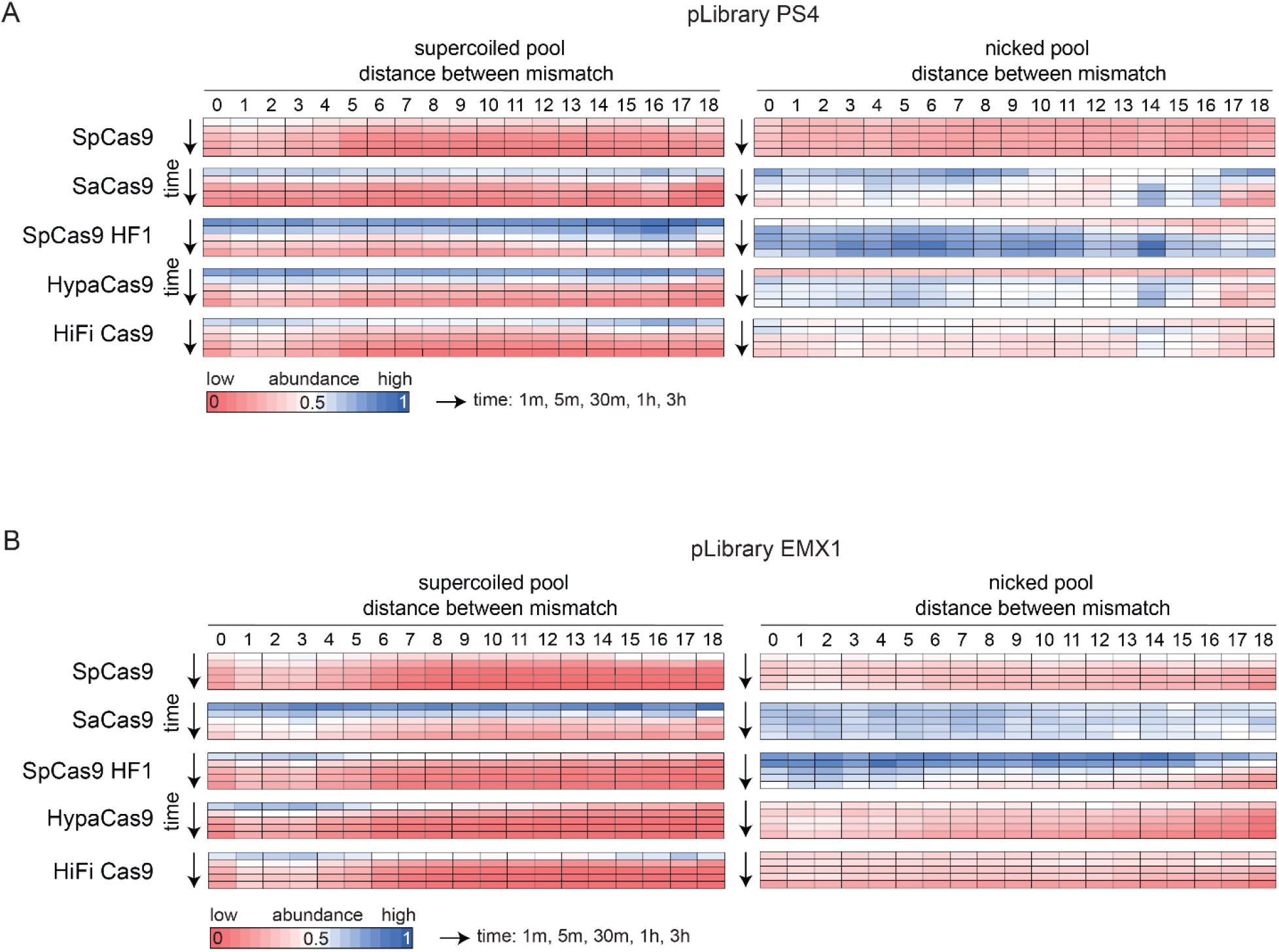
Effect of double mismatches in the target sequence on Cas9 cleavage activity. Heatmaps showing the relative abundance of target sequences with two mismatches over time for the supercoiled (left) and nicked (right) pools in pLibrary (A) PS4 and (B) EMX1 upon cleavage by Cas9 as a function of distance between the two mismatches and time. Time points indicated on the left by the arrow are 1 min, 5 min, 30 min, 1 h, and 3 h. Values plotted represent an average of two replicates.

We next looked at the positional effects when the two mismatches were present within the 10-nucloeotide PAM-proximal seed or the 10-nucleotide PAM-distal region. Here, the two mismatches could be separated by between 0 and 8 nucleotides. All Cas9s best tolerated mismatches that were separated by six or more nucleotides within the seed (Fig. S6A, B). The cleavage tolerance of two mismatches in the seed was similar among the HF Cas9 variants for pLibrary PS4 (Fig. S6A). However, there were notable differences for pLibrary EXM1 (Fig. S6B). While all Cas9 variants had defects for cleaving sequences with two mismatches separated by two to five nucleotides, SpCas9 HF1 and HypaCas9 depleted these sequences more rapidly from the supercoiled pool. In addition, all Cas9s except HypaCas9 accumulated sequences with two seed mismatches in the nicked pool (Fig. S6B). Together, these data suggest that HypaCas9 tolerated double mismatches in the seed for pLibrary EMX1 more than any other Cas9 variant.

Outside the seed region, most target sequences with two mismatches separated by any distance were eventually depleted from the supercoiled pool over time (Fig. S6C, D). For HF Cas9 variants, two mismatches separated by six or seven nucleotides in pLibrary PS4 were depleted more slowly from the supercoiled pool. Target sequences with two mismatches outside the seed separated by a distance of four nucleotides or less were more prone to nicking by SpCas9 HF1 for both pLibraries (Fig. S6C, D).

### Validating the nicking activity against mismatched targets

We observed an enrichment of target sequences with multiple mismatches in the nicked pool of the pLibraries upon Cas9 cleavage. We sought to validate this observation by verifying cleavage of target sequences present in the nicked pool of pLibrary PS4 subjected to cleavage by Cas9 at the longest time point (three hours). We tested cleavage activity of Cas9 variants against target sequences with two to five mismatches (MM). Similar to the HTS data, we observed varying degrees of nicking and linearization of target sequences containing different mismatches for different Cas9 variants (Fig. 7A). We quantified the supercoiled, linearized and nicked pools at two different time points, ten minutes and three hours for the perfect and mismatched target sequences (Fig. 7B). We observed that all Cas9s eventually fully linearized the perfect target (0 MM). Although the two and three mismatch target sequences tested contained two mismatches in the seed sequence, Cas9 linearized these targets with only about 20% remaining nicked after three hours of incubation. We observed little linearization and some nicking of the target sequences with seed mismatches, as in the case of pTarget 4.1 MM, indicating decreased cleavage activity of Cas9. However, Cas9 largely nicked target sequences with mismatches in the PAM-distal region, as in pTarget 4.2 MM. Interestingly, we also observed strong nicking of pTarget 5 MM by all Cas9s. Overall, all Cas9s showed more linearization activity against target sequences with up to three mismatches and nicking activity against targets with more mismatches.

**Figure 7.**
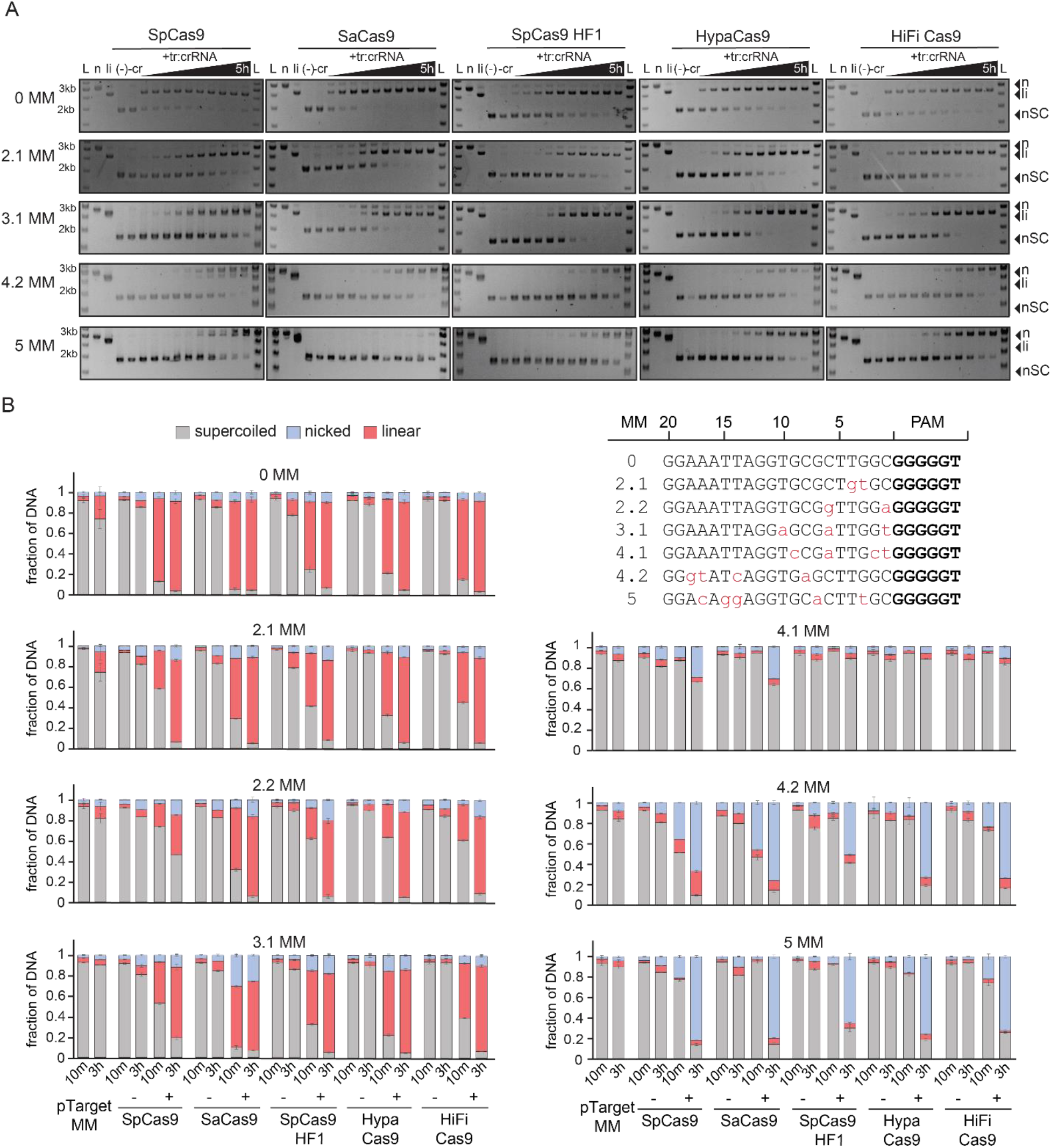
Cas9 variants have different cleavage activities against mismatched targets. (A) Representative agarose gels showing cleavage of a negatively supercoiled (nSC) plasmid containing the perfect target (0 MM) or mismatched (2 to 5 MM) target over a time course by Cas9 variants, resulting in linear (li) and/or nicked (n) products. Time points at which the samples were collected are 15 sec, 30 sec, 1 min, 2 min, 5 min, 15 min, 30 min, 1 h, 3 h, and 5 h. tr:crRNA = tracrRNA:crRNA. All controls were performed under the same conditions as the longest time point for the experimental samples. Controls: (−) = pTarget or pLibrary alone incubated at 37 °C for the longest time point in the assay (5 h); (-cr) = pTarget or pLibrary incubated with Cas9 only at 37°C for the longest time point in the assay (5 h); n = Nt.BspQI nicked pUC19; li = BsaI-HF linearized pUC19 (B) Quantification of supercoiled, linear and nicked pools from cleavage of perfect or fully crRNA-complementary (0 MM) and mismatched (2 to 5 MM) target plasmid by Cas9 after 10 minutes and 3 hours. pTarget MM indicates target plasmid (0, 2 to 5 MM) alone incubated at 37 °C for the time points indicated. (−) indicates a cleavage reaction with the target plasmid and Cas9 only, and (+) indicates a cleavage reaction with the target plasmid, Cas9 and cognate tracrRNA:crRNA. Values plotted represent an average of three replicates. Error bars are SEM. The different target sequences tested are listed where the PAM is in bold and mismatches are in lowercase and red.

## Discussion

Independent studies have reported the specificity of Cas9 orthologs and engineered variants separately (Chen et al., 2017; Hsu et al., 2013; Huston et al., 2019; Kleinstiver et al., 2016; Vakulskas et al., 2018; Zhang et al., 2020). Here, we directly compared the cleavage activity of five different Cas9 variants against two plasmid libraries. By studying the cleavage activities *in vitro*, we establish a comparative understanding of the native specificity of Cas9 variants. We show that Cas9 has target sequence-dependent linearization and nicking activities against targets with multiple mismatches. In agreement with Fu et al.’s *in vitro* studies (Fu et al., 2019, 2016, 2014), WT SpCas9 has sequence-dependent cleavage activity against targets with one or two mismatches. In addition, we establish that SpCas9 can nick and/or linearize sequences with up to five mismatches. This low-fidelity cleavage activity of SpCas9 has also been recorded in *in vivo* specificity studies (Fu et al., 2013; Hsu et al., 2013).

In parallel to SpCas9, we tested the cleavage activities of a natural ortholog, SaCas9 and three engineered SpCas9 variants – SpCas9 HF1, HypaCas9 and HiFi Cas9. Our analysis revealed that the high specificity of SaCas9 and HF Cas9 variants observed in previous studies may stem from the lack of detection of nicking by Cas9. Most studies test for DSB at target and off-target sites and/or indel formation (Chen et al., 2017; Huston et al., 2019; Kleinstiver et al., 2016; Schmid-Burgk et al., 2020; Vakulskas et al., 2018; Zhang et al., 2020). While SpCas9 linearized most target sequences with multiple mismatches as previously observed, SaCas9 and HF Cas9 variants often only nicked these sequences. Although HiFi Cas9 had similar on-target cleavage activity to SpCas9, it had higher off-target nicking activity.

Recent *in vitro* and *in vivo* studies profiled kinetic properties and genome editing activities of natural and high-fidelity Cas9 variants (Jones et al., 2019; Schmid-Burgk et al., 2020). In these studies, SpCas9 had high mismatch-tolerance and in turn, more off-target cleavage activity whereas HF Cas9 variants had reduced off-target cleavage activity. While the *in vitro* study concluded that SpCas9 HF1 had high cleavage specificity (reduced off-target cleavage rate compared to on-target cleavage), the *in vivo* study found that HiFi Cas9 had the best on-target activity and specificity ratio. Neither of the two studies were designed to address nicking of the target sequence. Nicked DNA may be subject to error-prone DNA repair, but nicks are more likely repaired by error-free DNA repair pathways (Fukui, 2010; Kuzminov, 2001; Vriend and Krawczyk, 2017). It is also important to note that off-targets that can be nicked by Cas9 may also be sequestered by chromatin structure in vivo (Horlbeck et al., 2016; Yarrington et al., 2018). Nevertheless, new methods developed to detect nicks may be used to detect potential off-target nicking activity that are unreported in the previous studies (Cao et al., 2019; Elacqua et al., 2019).

Cas9 cleavage specificity depends on both the position-specific mismatches (within or outside the seed) and the target sequence (Hsu et al., 2013; Huston et al., 2019; Liu et al., 2016; Zhang et al., 2020). We tested two different pLibraries that show the effects of target-crRNA sequence and various nucleotide substitutions across the target region. Our heatmaps show that mismatches in the PAM-distal region of the target sequence in both pLibraries contribute to target nicking rather than DSBs. While SpCas9 had a similar specificity score for both pLibraries, the specificity of other Cas9 variants not only depended on the target sequence but also the time of exposure. SaCas9 had a similar specificity score as SpCas9 for both pLibraries despite the differences in the cleavage activities against these pLibraries. SpCas9 HF1 and HypaCas9 had similar specificity scores as SpCas9 at longer incubation times. HiFi Cas9, however had a similar cleavage activity against both pLibraries tested but was more specific compared to SpCas9.

Cas9 undergoes conformational changes in the REC domain upon binding to its guide RNA (Anders et al., 2014; Jinek et al., 2014; Nishimasu et al., 2014). Following target recognition and binding, further domain rearrangements occur. Cas9 cleavage activity is limited by R-loop formation upon target recognition (Gong et al., 2018; Singh et al., 2018). HF Cas9 variants were reported to have higher target sequence unwinding specificity compared to SpCas9 (Okafor et al., 2019). HF Cas9 variants can also discriminate against cleavage of target sequences with mismatches due to decreased rates of cleavage, which may result in off-target DNA release rather than cleavage (Liu et al., 2019). SpCas9 HF1 was designed to disrupt contacts made by Cas9 and the target strand (Kleinstiver et al., 2016). This alters target binding with a stringent requirement of base-pairing between the crRNA and target DNA. This possibly leads to R-loop destabilization and in turn, discrimination against targets with mismatches for cleavage as seen in *in vivo* studies. *In vitro* binding and cleavage analysis of an oligonucleotide library by HiFi Cas9 showed reduced cleavage of target sequences with mismatches compared to SpCas9 (Zhang et al., 2020). Our pLibrary cleavage assays indeed showed slower rates of cleavage of target sequences with mismatches by HF Cas9 variants. Although, not all target sequences with mismatches were fully cleaved by HF Cas9 variants compared to SpCas9, the reduced rate of linearization and nicking may possibly be due to R-loop collapse or premature release of these targets following nicking.

Cas9 cleaves dsDNA using the HNH and RuvC domains that cleave the crRNA-complementary target strand and non-target strand, respectively (Chen et al., 2014; Jinek et al., 2012). Upon binding to a target, the HNH domain is re-positioned for target strand cleavage. However, some mismatches in the target sequence may prevent HNH domain movement (Dagdas et al., 2017; Sternberg et al., 2015). RuvC domain activation is allosterically controlled by HNH conformational changes but HNH nuclease activity is not a prerequisite. HypaCas9 was designed with mutations in the REC domain that prevent HNH domain activation and movement (Chen et al., 2017). Both SpCas9 HF1 and HiFi Cas9 have a single mutation in the REC domain (Kleinstiver et al., 2016; Vakulskas et al., 2018). Upon binding to a target sequence with mismatches, sufficient HNH domain movement may trigger cleavage of the non-target strand by the RuvC domain without activation of the HNH domain for target strand cleavage, leading to nicking of the non-target strand.

Cas9 generates DSB in the target with little post-cleavage trimming of the non-target strand (Jinek et al., 2012; Stephenson et al., 2018). Unlike other CRISPR systems where a Cas nuclease has processive cleavage activity to clear the invading phage nucleic acid upon recognition and/or cleavage, Cas9-containing host bacteria may rely on host nucleases to clear the Cas9-targeted phage DNA (Hille et al., 2018). Despite the lack of any strong post-cleavage activities, Cas9 provides robust protection against phages (Barrangou et al., 2007). This is possibly due to the high mismatch tolerance of Cas9 which may enable better protection against closely related and rapidly evolving phages. Additionally, the ability to nick target sequences with multiple mismatches may also contribute to the immunity against phages. Off-target, sequence-dependent nicking or non-specific nicking have been recently reported for several other Cas effector proteins (McMahon et al., 2020; Murugan et al., 2020; Yan et al., 2019). Here, target sequence-dependent nicking activity may help to slow down phage replication and fight against phages that develop mutations in the target region (McMahon et al., 2020; Tao et al., 2018). Further studies are required to demonstrate the benefits of target sequence-dependent nicking activity *in vivo*.

## Methods

### Cas9 expression vectors

Expression plasmids for SpCas9 and high-fidelity variants were purchased from Addgene. *Streptococcus pyogenes* Cas9 (SpCas9) (pMJ806) was expressed using expression plasmid pEC-K-MBP, and SpCas9-HF1 (pJSC111) and HypaCas9 (pJSC173) were expressed using expression plasmid pCT10. pMJ806, pJSC111, pJSC173 were gifts from Jennifer Doudna and/or Keith Joung (Addgene plasmid # 39312; http://n2t.net/addgene:39312; RRID:Addgene_39312; Addgene plasmid # 101209; http://n2t.net/addgene:101209; RRID:Addgene_101209; Addgene plasmid # 101218; http://n2t.net/addgene:101218; RRID:Addgene_101218). The gene sequence for *Staphylococcus aureus* Cas9 (SaCas9) was synthesized as *Escherichia coli* codon-optimized gBlocks (purchased from Integrated DNA Technologies, IDT). SaCas9 gBlocks were cloned into pSV272 with 6X-His sequence, a maltose binding protein (MBP) and a Tobacco Etch Virus (TEV) protease cleavage site in the N-terminal via Gibson assembly (New England Biolabs) as per the manufacturer’s protocol.

### Cas9 expression and purification

All Cas9 proteins were expressed in *Escherichia coli* BL21 (DE3) cells. Overnight cultures of the cells carrying the expression plasmid was used to inoculate 2X TY broth supplemented with corresponding antibiotics in 1:100 ratio. Cultures were grown at 37 °C to an optical density (600 nm) of 0.5 − 0.6 and IPTG was added to a final concentration of 0.2 mM to induce protein expression. The incubation was continued at 18 °C overnight (~16 – 18 hours) and harvested the next day for protein purification.

SpCas9 was purified by the following protocol (Chen et al., 2014). Cells were resuspended in Lysis Buffer I (20 mM Tris-HCl pH 8.0, 500 mM NaCl, 10 mM imidazole, and 10% glycerol) supplemented with PMSF. A sonicator or homogenizer was used to lyse the cells and the lysate was centrifuged to remove insoluble material. The clarified lysate was applied to a HisPur™ Ni-NTA Resin (ThermoFisher Scientific) column. After washing the column with Lysis Buffer I, the bound protein was eluted in Elution Buffer I (Lysis Buffer I + 250 mM imidazole final concentration). The Ni-NTA column eluent was concentrated and run on a HiLoad 16/600 Superdex 200 gel filtration column (GE Healthcare) pre-equilibrated with SEC Buffer A (20 mM Tris-HCl, pH 8.0, and 500 mM NaCl). TEV protease was added at 1:100 (w/w) ratio to the pools containing 6X His-MBP tagged Cas9 and left on ice, overnight at 4 °C. Samples were reapplied to HisPur™ Ni-NTA Resin (ThermoFisher Scientific) to remove the His-tagged TEV, free 6X His-MBP, and any remaining tagged protein. The flow-through was collected, concentrated and further purified by using a HiLoad 16/600 S200 gel filtration column in SEC Buffer B (20 mM Tris-HCl, pH 8.0, 200 mM KCl, and 1mM EDTA). Peak pools were analyzed on SDS-PAGE gels and the pools with Cas9 were combined, concentrated, flash frozen in liquid nitrogen and stored at −80°C until further use.

The purification protocol was further optimized and all other Cas9 variants were purified using the following protocol (Mohanraju et al., 2018). Harvested cells were resuspended in Lysis Buffer II (20 mM Tris-HCl pH 8.0, 500 mM NaCl, 5 mM imidazole), supplemented with protease inhibitors (PMSF, cOmplete™ Protease Inhibitor Cocktail Tablet or Halt Protease Inhibitor Cocktail). A sonicator or homogenizer was used to lyse the cells and the lysate was centrifuged to remove insoluble material. The clarified lysate was applied to a HisPur™ Ni-NTA Resin (ThermoFisher Scientific) column. After washing the column with 10 column volumes of Wash Buffer (Lysis Buffer + 15 mM imidazole final concentration), the bound protein was eluted in Elution Buffer I (Lysis Buffer II + 250 mM imidazole final concentration). Fractions containing Cas9 were pooled and TEV protease was added in a 1:100 (w/w) ratio and dialyzed in Dialysis Buffer (10 mM HEPES-KOH pH 7.5, 200 mM KCl, 1 mM DTT) at 4°C overnight. The dialyzed protein was diluted 1:1 with 20 mM HEPES KOH (pH 7.5) and loaded on a HiTrap Heparin HP (GE Healthcare) column and washed with Buffer A (20 mM HEPES-KOH pH 7.5, 100 mM KCl). The protein was eluted with Buffer B (20 mM HEPES-KOH pH 7.5, 2 M KCl) by applying a gradient from 0% to 50% over a total volume of 60 ml. Eluted peak fractions were analyzed by SDS-PAGE and fractions with Cas9 were combined and concentrated. DTT was added to a final concentration of 1 mM. The protein was fractionated on a HiLoad 16/600 Superdex 200 gel filtration column (GE Healthcare), eluting with SEC buffer (20 mM HEPES-KOH pH 7.5, 500 mM KCl, 1mM DTT). Peak pools were analyzed on SDS-PAGE gels and the pools with Cas9 were combined, concentrated, flash frozen in liquid nitrogen and stored at −80°C until further use.

Alt-R ® S.p. HiFi Cas9 was provided by Integrated DNA Technologies (IDT) – HiFi Cas9.

### Library creation

As previously described (Murugan et al., 2020), the library was partially randomized to generate a pool of sequences containing mismatches (Pollard et al., 2000). The following probability distribution function was used to determine the randomization/doping frequency

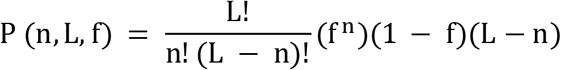

where, P is the pool of the population, L is the sequence length, n is the number of mutations/template and f is the probability of mutation/position (doping level or frequency). A randomization/doping frequency (f) of 15% was selected to optimize the library to contain a maximum of sequences with 2 to 4 mismatches.

The number of different mutation combinations (MM_c_) for a given number of mutations, n and sequence length, L, regardless of the doping level/frequency is determined by,

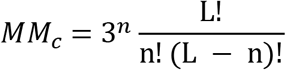

The total number of unique target sequences with a single mismatch is 60, with 2 mismatches is 1,710, and with 3 mismatches is 30,780, etc. We used two library sequences that we previously tested (Murugan et al., 2020), a modified protospacer 4 sequence from *Streptococcus pyogenes* CRISPR locus (55% GC) and EMX1 gene target sequence (80% GC) (see Supplementary Table 1 for target sequence).

### Plasmid and nucleic acid preparation

All DNA oligonucleotides used in this study were synthesized by IDT or Thermo Scientific. RNAs (tracrRNA and crRNA) and single-stranded library oligonucleotides with 15% doping frequency in the target region were ordered from IDT. Supplementary Table 1 has the sequences of DNA oligonucleotides and RNA used in this study.

Gibson assembly was used to generate libraries and target plasmids. The oligonucleotides for the libraries, target and mismatched targets were diluted to 0.2 μM in 1X NEBuffer 2. pUC19 vector was amplified using primers listed in supplementary table 1 via PCR to insert homology arms. The PCR reaction was subjected to DpnI digestion and PCR cleanup (Promega Wizard SV Gel and PCR Clean-Up System). 30 ng of PCR amplified pUC19, 5 μL of oligonucleotide (0.2 μM) and ddH2O to bring the volume to 10 μL were mixed with 10 μL 2X NEBuilder HiFi DNA Assembly Master mix (New England Biolabs) and incubated at 50 °C for 1 hour. NEB Stable competent cells were transformed with 2 μL of the assembled product, as per the manufacturer’s protocol. After the recovery step, all of cells in the outgrowth media were used to inoculate 50 mL LB supplemented with ampicillin and incubated overnight at 37 °C. The following precautions were taken to ensure the plasmid was intact and remain supercoiled after extractions as described before (Murugan et al., 2020). Cells were cooled on ice before harvesting for the plasmid extraction using QIAGEN Plasmid Midi Kit. All the initial steps from lysis to neutralization for plasmid extractions (pTarget, pLibrary and empty plasmid) were performed on ice with minimum mechanical stress. Plasmid were stored as aliquots that were used for up to 10 freeze-thaw cycles. Two different pLibrary assembly reactions and preparations were used for the two replicates of the high-throughput *in vitro* cleavage assays (Fig. S2A).

For controls, target plasmids and empty pUC19 were linearized by restriction enzyme digestion using BsaI-HF and nicked using a nicking enzyme Nt.BspQI (New England Biolabs). All restriction digestion reactions were carried out as per the manufacturer’s protocols. All sequences were verified by Sanger sequencing (Eurofins Genomics, Kentucky, USA). The topology of the extracted and restriction digested plasmids was verified on an agarose gel before using in cleavage assays (Fig. S2A).

### In vitro cleavage assay and analysis

The protocol was adapted from previously described methods (Anders and Jinek, 2014). Briefly, Cas9:tracrRNA:crRNA complex was formed by incubating Cas9 and tracrRNA:crRNA (1:1.5 ratio) in 1X reaction buffer (20 mM HEPES, pH 7.4, 100 mM potassium chloride, 5 mM magnesium chloride, 1 mM dithiothreitol, and 5% glycerol) at 37 °C for 10 min. Cas9 RNP complex was mixed with pTarget, pLibrary or empty plasmid (150 ng) to initiate cleavage reactions and incubating at 37 °C. Phenol-chloroform was used to quench reactions at different time points. The aqueous layer was extracted and separated on a 1% agarose gel via electrophoresis and stained with SYBR safe or RED safe stain for dsDNA visualization. Excess tracrRNA:crRNA was used in cleavage assays to prevent any RNA-independent cleavage activity (Sundaresan et al., 2017). For library, mismatched target plasmid and empty plasmid cleavage assays, 100 nM Cas9 and 150 nM tracrRNA:crRNA was used. Concentrations of pLibrary, pTarget and empty pUC19 used were at 150 ng/10 μL (8.6 nM) reaction.

Bands were visualized and quantified with ImageJ (https://imagej.nih.gov/ij/). Intensities of the band (I) in the uncleaved (supercoiled-SC) and cleaved fractions (nicked-N and linearized-L) were measured. Fractions (FR) cleaved and uncleaved were calculated as follows.

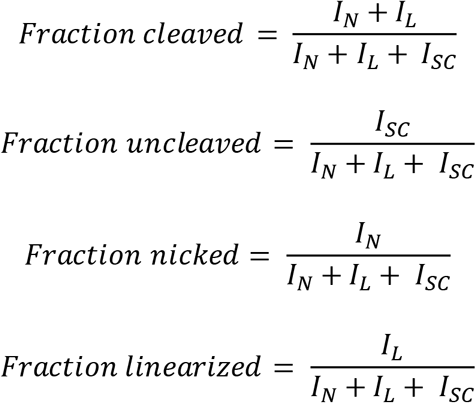

The FR_SC_, FR_N_ and FR_L_ were determined for each of the time points ‘t’. FR for time point 0 was the determined for the negative control pLibrary (i.e. pLibrary run on a gel after preparation as represented in Fig. S2A).

### Library preparation for HTS

Agarose gel electrophoresis (as described above) was used to separate the library plasmid cleavage products into cleaved (linear and nicked) and uncleaved (supercoiled) products. The bands from the nicked and supercoiled pools from various time points were excised separately and were individually gel purified using QIAquick Gel Extraction Kit (Qiagen). Nextera Adapters (NEA) designed to amplify the target region in the pLibrary and standard Nextera unique indices/barcodes to multiplex the samples were added to with two rounds of PCRs (see Supplementary Table 1 for NEA primers). Samples were cleaned using QIAquick PCR Purification Kit (Qiagen) between the two PCR steps. The size of the PCR products was verified using Agilent 2100 Bioanalyzer. Pooled samples were subjected NextSeq or MiSeq for paired-end reads of 75 cycles at Admera Health, LLC (New Jersey, USA) or Iowa State DNA Facility (Ames, IA). 15% PhiX was spiked in.

### HTS data analysis

The analysis of HTS data was done as previously described (Murugan et al., 2020). Briefly, the library HTS data were processed with custom bash scripts (see associated GitHub repository: https://github.com/sashital-lab/Cas9_specificity). A simple workflow of the analysis is described in supplementary figure 1, adapted from our previous study on Cas12a (Murugan et al., 2020). Target sequences were extract along with the counts of the extracted target sequences and the number of mismatches. Target sequence information were imported into Microsoft Excel for plotting and summarizing, post command-line processing.

In each pool, the fraction of target sequences containing ‘n’ mismatches (MM) (F_n-MM_) was calculated as follows.

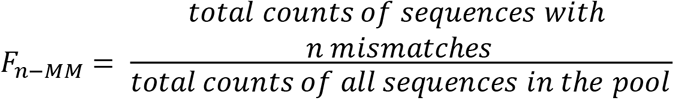

FR was calculated for each time point as described above in *In vitro* cleavage assay and analysis section. F_n-MM_ was normalized to the fraction (FR) of DNA present in the supercoiled or nicked fraction at a given time point ‘t’ to generate an estimated abundance (EA) of a given set of sequences at a given timepoint.

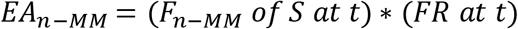

These values were plotted against number of mismatches to generate mismatch distribution curves. The relative abundance (enrichment and/or depletion) (RA) of a sequence ‘S’ containing ‘n’ mismatches at each time point ‘t’ compared to the pLibraries in the negative control, (i.e. pLibrary run on a gel after preparation as represented in Fig. S2A) was calculated and plotted versus time to generate normalized curves.

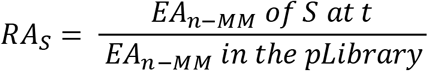

The RA for the perfect target sequence (0 MM), RA_0MM_ was calculated using the above equation. The RA for target sequences with 1 to 5 MM at each time point ‘t’, RA_1-5MM-t_ was calculated by summing EA for 1 to 5 MM, EA_1-5MM-t_ and normalizing to the sum of EA of 1 to 5 MM in the negative control pLibrary, EA_1-5MM-0_ (i.e. pLibrary run on a gel after preparation as represented in Fig. S2A). The relative cleaved fraction of counts or abundance (RA_FR_) was determined by subtracting these RA_0MM_ or RA_1-5MM_ values from 1 and plotted against time. The specificity score for Cas9 cleavage was calculated by dividing RA_0MM_ by RA_1-5MM_ (RA_0MM_/ RA_1-5MM_).

For the heatmaps, the estimated abundance (EA) of sequences containing a particular nucleotide (N = A, G, C, T) at a particular position (P = 1 to 20) for target sequences containing ‘n’ mismatches at each time point ‘t’ was calculated as above. Relative abundance (RA) was calculated by normalizing EA against the pool of DNA in the original library to eliminate variability in aberrant nicking that may have occurred for individual pLibraries in the negative control.

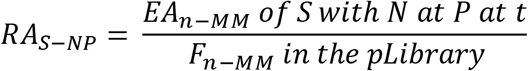

The cleavage rates differed among the Cas9 variants and against target sequences with varying mismatches. Therefore, each RA value was normalized to the maximum RA value present within each Cas9 and mismatch set in either the supercoiled or nicked pool to scale the relative abundance from 0 to 1 for the heatmaps. For the analysis of target sequences with two mismatches, the sequences with 2 mismatches were extracted. The distance between the two mismatches and the total counts for sequences separated by that distance were determined. The counts were normalized to the number of possible ways the two mismatches can occur, and the normalized RA was calculated similarly to the heatmaps.

## Acknowledgements

We thank all the former and current members of the Sashital Lab for helpful discussions and suggestions on various aspects of the project. We would like to thank Mollie Schubert and Aftan Vander Zwaag at Integrated DNA Technologies, Inc. (IDT), Coralville IA for providing initial samples of Alt-R ® S.p. HiFi Cas9. We also thank Heather S. Lewin and Megan N. O’Donnell from the University Library for helping with the data deposition to DataShare. This work was supported by startup funds to D.G.S from Iowa State University College of Liberal Arts and Sciences and the Roy J. Carver Charitable Trust, the National Science Foundation (1652661 to D.G.S), and National Institute of Food and Agriculture (IOW05480 to D.G.S).

## Author contributions

All experiments and HTS sample preparation, were performed by K.M. HTS data extraction was performed by A.J.S using bash scripts written by A.S.S. HTS results were analyzed and interpreted by K.M and D.G.S. The manuscript was written by K.M and D.G.S, with inputs from A.J.S and A.S.S. Funding for this project was secured by D.G.S.

## Data Availability

HTS data and processed data files from this study have been deposited in the Iowa State University Library’s DataShare, and can be found at https://doi.org/10.25380/iastate.12245846. All other information and data are available from the authors upon request.

## References

Amrani, N., Gao, X.D., Liu, P., Edraki, A., Mir, A., Ibraheim, R., Gupta, A., Sasaki, K.E., Wu, T., Donohoue, P.D., Settle, A.H., Lied, A.M., McGovern, K., Fuller, C.K., Cameron, P., Fazzio, T.G., Zhu, L.J., Wolfe, S.A., Sontheimer, E.J., 2018. NmeCas9 is an intrinsically high-fidelity genome-editing platform. Genome Biology 19, 214. https://doi.org/10.1186/s13059-018-1591-1

Anders, C., Jinek, M., 2014. In Vitro Enzymology of Cas9, in: Methods in Enzymology. Elsevier, pp. 1–20. https://doi.org/10.1016/B978-0-12-801185-0.00001-5

Anders, C., Niewoehner, O., Duerst, A., Jinek, M., 2014. Structural basis of PAM-dependent target DNA recognition by the Cas9 endonuclease. Nature 513, 569–573. https://doi.org/10.1038/nature13579

Barrangou, R., Fremaux, C., Deveau, H., Richards, M., Boyaval, P., Moineau, S., Romero, D.A., Horvath, P., 2007. CRISPR Provides Acquired Resistance Against Viruses in Prokaryotes. Science 315, 1709–1712. https://doi.org/10.1126/science.1138140

Cao, B., Wu, X., Zhou, J., Wu, H., DeMott, M.S., Gu, C., Wang, L., You, D., Dedon, P.C., 2019. Nick-seq for single-nucleotide resolution genomic maps of DNA modifications and damage (preprint). Molecular Biology. https://doi.org/10.1101/845768

Chen, H., Choi, J., Bailey, S., 2014. Cut Site Selection by the Two Nuclease Domains of the Cas9 RNA-guided Endonuclease. J Biol Chem 289, 13284–13294. https://doi.org/10.1074/jbc.M113.539726

Chen, J.S., Dagdas, Y.S., Kleinstiver, B.P., Welch, M.M., Sousa, A.A., Harrington, L.B., Sternberg, S.H., Joung, J.K., Yildiz, A., Doudna, J.A., 2017. Enhanced proofreading governs CRISPR–Cas9 targeting accuracy. Nature 550, 407–410. https://doi.org/10.1038/nature24268

Cong, L., Ran, F.A., Cox, D., Lin, S., Barretto, R., Habib, N., Hsu, P.D., Wu, X., Jiang, W., Marraffini, L.A., Zhang, F., 2013. Multiplex Genome Engineering Using CRISPR/Cas Systems. Science 339, 819–823. https://doi.org/10.1126/science.1231143

Dagdas, Y.S., Chen, J.S., Sternberg, S.H., Doudna, J.A., Yildiz, A., 2017. A conformational checkpoint between DNA binding and cleavage by CRISPR-Cas9. Science Advances 3, eaao0027. https://doi.org/10.1126/sciadv.aao0027

Doudna, J.A., 2020. The promise and challenge of therapeutic genome editing. Nature 578, 229–236. https://doi.org/10.1038/s41586-020-1978-5

Elacqua, J.J., Ranu, N., Dilorio, S.E., Blainey, P.C., 2019. NickSeq for genome-wide strand-specific identification of DNA single-strand break sites with single nucleotide resolution (preprint). Genomics. https://doi.org/10.1101/867937

Friedland, A.E., Baral, R., Singhal, P., Loveluck, K., Shen, S., Sanchez, M., Marco, E., Gotta, G.M., Maeder, M.L., Kennedy, E.M., Kornepati, A.V.R., Sousa, A., Collins, M.A., Jayaram, H., Cullen, B.R., Bumcrot, D., 2015. Characterization of Staphylococcus aureus Cas9: a smaller Cas9 for all-in-one adeno-associated virus delivery and paired nickase applications. Genome Biology 16, 257. https://doi.org/10.1186/s13059-015-0817-8

Fu, B.X.H., Hansen, L.L., Artiles, K.L., Nonet, M.L., Fire, A.Z., 2014. Landscape of target:guide homology effects on Cas9-mediated cleavage. Nucleic Acids Res 42, 13778–13787. https://doi.org/10.1093/nar/gku1102

Fu, B.X.H., Smith, J.D., Fuchs, R.T., Mabuchi, M., Curcuru, J., Robb, G.B., Fire, A.Z., 2019. Target-dependent nickase activities of the CRISPR–Cas nucleases Cpf1 and Cas9. Nature Microbiology. https://doi.org/10.1038/s41564-019-0382-0

Fu, B.X.H., St. Onge, R.P., Fire, A.Z., Smith, J.D., 2016. Distinct patterns of Cas9 mismatch tolerance in vitro and in vivo. Nucleic Acids Res 44, 5365–5377. https://doi.org/10.1093/nar/gkw417

Fu, Y., Foden, J.A., Khayter, C., Maeder, M.L., Reyon, D., Joung, J.K., Sander, J.D., 2013. High-frequency off-target mutagenesis induced by CRISPR-Cas nucleases in human cells. Nature Biotechnology 31, 822–826. https://doi.org/10.1038/nbt.2623

Fukui, K., 2010. DNA Mismatch Repair in Eukaryotes and Bacteria [WWW Document]. Journal of Nucleic Acids. https://doi.org/10.4061/2010/260512

Gasiunas, G., Barrangou, R., Horvath, P., Siksnys, V., 2012. Cas9–crRNA ribonucleoprotein complex mediates specific DNA cleavage for adaptive immunity in bacteria. PNAS 109, E2579–E2586. https://doi.org/10.1073/pnas.1208507109

Gong, S., Yu, H.H., Johnson, K.A., Taylor, D.W., 2018. DNA Unwinding Is the Primary Determinant of CRISPR-Cas9 Activity. Cell Reports 22, 359–371. https://doi.org/10.1016/j.celrep.2017.12.041

Hille, F., Richter, H., Wong, S.P., Bratovič, M., Ressel, S., Charpentier, E., 2018. The Biology of CRISPR-Cas: Backward and Forward. Cell 172, 1239–1259. https://doi.org/10.1016/j.cell.2017.11.032

Horlbeck, M.A., Witkowsky, L.B., Guglielmi, B., Replogle, J.M., Gilbert, L.A., Villalta, J.E., Torigoe, S.E., Tjian, R., Weissman, J.S., 2016. Nucleosomes impede Cas9 access to DNA in vivo and in vitro. eLife 5, e12677. https://doi.org/10.7554/eLife.12677

Hsu, P.D., Scott, D.A., Weinstein, J.A., Ran, F.A., Konermann, S., Agarwala, V., Li, Y., Fine, E.J., Wu, X., Shalem, O., Cradick, T.J., Marraffini, L.A., Bao, G., Zhang, F., 2013. DNA targeting specificity of RNA-guided Cas9 nucleases. Nature Biotechnology 31, 827–832. https://doi.org/10.1038/nbt.2647

Hu, J.H., Miller, S.M., Geurts, M.H., Tang, W., Chen, L., Sun, N., Zeina, C.M., Gao, X., Rees, H.A., Lin, Z., Liu, D.R., 2018. Evolved Cas9 variants with broad PAM compatibility and high DNA specificity. Nature 556, 57–63. https://doi.org/10.1038/nature26155

Huston, N.C., Tycko, J., Tillotson, E.L., Wilson, C.J., Myer, V.E., Jayaram, H., Steinberg, B.E., 2019. Identification of Guide-Intrinsic Determinants of Cas9 Specificity. The CRISPR Journal 2, 172–185. https://doi.org/10.1089/crispr.2019.0009

Jiang, F., Doudna, J.A., 2017. CRISPR–Cas9 Structures and Mechanisms. Annual Review of Biophysics 46, 505–529. https://doi.org/10.1146/annurev-biophys-062215-010822

Jinek, M., Chylinski, K., Fonfara, I., Hauer, M., Doudna, J.A., Charpentier, E., 2012. A Programmable Dual-RNA–Guided DNA Endonuclease in Adaptive Bacterial Immunity. Science 337, 816–821. https://doi.org/10.1126/science.1225829

Jinek, M., Jiang, F., Taylor, D.W., Sternberg, S.H., Kaya, E., Ma, E., Anders, C., Hauer, M., Zhou, K., Lin, S., Kaplan, M., Iavarone, A.T., Charpentier, E., Nogales, E., Doudna, J.A., 2014. Structures of Cas9 Endonucleases Reveal RNA-Mediated Conformational Activation. Science 343. https://doi.org/10.1126/science.1247997

Jones, S.K., Hawkins, J.A., Johnson, N.V., Jung, C., Hu, K., Rybarski, J.R., Chen, J.S., Doudna, J.A., Press, W.H., Finkelstein, I.J., 2019. Massively parallel kinetic profiling of natural and engineered CRISPR nucleases. bioRxiv 696393. https://doi.org/10.1101/696393

Kim, E., Koo, T., Park, S.W., Kim, D., Kim, K., Cho, H.-Y., Song, D.W., Lee, K.J., Jung, M.H., Kim, S., Kim, Jin Hyoung, Kim, Jeong Hun, Kim, J.-S., 2017. In vivo genome editing with a small Cas9 orthologue derived from Campylobacter jejuni. Nature Communications 8, 1–12. https://doi.org/10.1038/ncomms14500

Kleinstiver, B.P., Pattanayak, V., Prew, M.S., Tsai, S.Q., Nguyen, N.T., Zheng, Z., Joung, J.K., 2016. High-fidelity CRISPR–Cas9 nucleases with no detectable genome-wide off-target effects. Nature 529, 490–495. https://doi.org/10.1038/nature16526

Kuzminov, A., 2001. Single-strand interruptions in replicating chromosomes cause double-strand breaks. PNAS 98, 8241–8246. https://doi.org/10.1073/pnas.131009198

Lee, J.K., Jeong, E., Lee, J., Jung, M., Shin, E., Kim, Y., Lee, K., Jung, I., Kim, D., Kim, S., Kim, J.-S., 2018. Directed evolution of CRISPR-Cas9 to increase its specificity. Nature Communications 9, 1–10. https://doi.org/10.1038/s41467-018-05477-x

Liu, M.-S., Gong, S., Yu, H.-H., Jung, K., Johnson, K.A., Taylor, D.W., 2019. Basis for discrimination by engineered CRISPR/Cas9 enzymes. bioRxiv 630509. https://doi.org/10.1101/630509

Liu, X., Homma, A., Sayadi, J., Yang, S., Ohashi, J., Takumi, T., 2016. Sequence features associated with the cleavage efficiency of CRISPR/Cas9 system. Scientific Reports 6, 19675. https://doi.org/10.1038/srep19675

Makarova, K.S., Wolf, Y.I., Iranzo, J., Shmakov, S.A., Alkhnbashi, O.S., Brouns, S.J.J., Charpentier, E., Cheng, D., Haft, D.H., Horvath, P., Moineau, S., Mojica, F.J.M., Scott, D., Shah, S.A., Siksnys, V., Terns, M.P., Venclovas, Č., White, M.F., Yakunin, A.F., Yan, W., Zhang, F., Garrett, R.A., Backofen, R., van der Oost, J., Barrangou, R., Koonin, E.V., 2020. Evolutionary classification of CRISPR–Cas systems: a burst of class 2 and derived variants. Nature Reviews Microbiology 18, 67–83. https://doi.org/10.1038/s41579-019-0299-x

Mali, P., Yang, L., Esvelt, K.M., Aach, J., Guell, M., DiCarlo, J.E., Norville, J.E., Church, G.M., 2013. RNA-Guided Human Genome Engineering via Cas9. Science 339, 823–826. https://doi.org/10.1126/science.1232033

McMahon, S.A., Zhu, W., Graham, S., Rambo, R., White, M.F., Gloster, T.M., 2020. Structure and mechanism of a Type III CRISPR defence DNA nuclease activated by cyclic oligoadenylate. Nature Communications 11, 1–11. https://doi.org/10.1038/s41467-019-14222-x

Mohanraju, P., Oost, J., Jinek, M., Swarts, D., 2018. Heterologous Expression and Purification of the CRISPR-Cas12a/Cpf1 Protein. BIO-PROTOCOL 8. https://doi.org/10.21769/BioProtoc.2842

Murugan, K., Seetharam, A.S., Severin, A.J., Sashital, D.G., 2020. CRISPR-Cas12a has widespread off-target and dsDNA-nicking effects. J. Biol. Chem. jbc.RA120.012933. https://doi.org/10.1074/jbc.RA120.012933

Nishimasu, H., Ran, F.A., Hsu, P.D., Konermann, S., Shehata, S.I., Dohmae, N., Ishitani, R., Zhang, F., Nureki, O., 2014. Crystal Structure of Cas9 in Complex with Guide RNA and Target DNA. Cell 156, 935–949. https://doi.org/10.1016/j.cell.2014.02.001

Okafor, I.C., Singh, D., Wang, Y., Jung, M., Wang, H., Mallon, J., Bailey, S., Lee, J.K., Ha, T., 2019. Single molecule analysis of effects of non-canonical guide RNAs and specificity-enhancing mutations on Cas9-induced DNA unwinding. Nucleic Acids Res 47, 11880–11888. https://doi.org/10.1093/nar/gkz1058

Oppenheim, A., 1981. Separation of closed circular DNA from linear DNA by electrophoresis in two dimensions in agarose gels. Nucleic Acids Res 9, 6805–6812. https://doi.org/10.1093/nar/9.24.6805

Pattanayak, V., Lin, S., Guilinger, J.P., Ma, E., Doudna, J.A., Liu, D.R., 2013. High-throughput profiling of off-target DNA cleavage reveals RNA-programmed Cas9 nuclease specificity. Nature Biotechnology 31, 839–843. https://doi.org/10.1038/nbt.2673

Pollard, J., Bell, S.D., Ellington, A.D., 2000. Design, Synthesis, and Amplification of DNA Pools for In Vitro Selection. Current Protocols in Nucleic Acid Chemistry 00, 9.2.1–9.2.23. https://doi.org/10.1002/0471142700.nc0902s00

Ran, F.A., Cong, L., Yan, W.X., Scott, D.A., Gootenberg, J.S., Kriz, A.J., Zetsche, B., Shalem, O., Wu, X., Makarova, K.S., Koonin, E.V., Sharp, P.A., Zhang, F., 2015. In vivo genome editing using Staphylococcus aureus Cas9. Nature 520, 186–191. https://doi.org/10.1038/nature14299

Schmid-Burgk, J.L., Gao, L., Li, D., Gardner, Z., Strecker, J., Lash, B., Zhang, F., 2020. Highly Parallel Profiling of Cas9 Variant Specificity. Molecular Cell. https://doi.org/10.1016/j.molcel.2020.02.023

Singh, D., Wang, Y., Mallon, J., Yang, O., Fei, J., Poddar, A., Ceylan, D., Bailey, S., Ha, T., 2018. Mechanisms of improved specificity of engineered Cas9s revealed by single-molecule FRET analysis. Nature Structural & Molecular Biology 25, 347–354. https://doi.org/10.1038/s41594-018-0051-7

Slaymaker, I.M., Gao, L., Zetsche, B., Scott, D.A., Yan, W.X., Zhang, F., 2016. Rationally engineered Cas9 nucleases with improved specificity. Science 351, 84–88. https://doi.org/10.1126/science.aad5227

Stephenson, A.A., Raper, A.T., Suo, Z., 2018. Bidirectional Degradation of DNA Cleavage Products Catalyzed by CRISPR/Cas9. J. Am. Chem. Soc. 140, 3743–3750. https://doi.org/10.1021/jacs.7b13050

Sternberg, S.H., LaFrance, B., Kaplan, M., Doudna, J.A., 2015. Conformational control of DNA target cleavage by CRISPR–Cas9. Nature 527, 110–113. https://doi.org/10.1038/nature15544

Sternberg, S.H., Redding, S., Jinek, M., Greene, E.C., Doudna, J.A., 2014. DNA interrogation by the CRISPR RNA-guided endonuclease Cas9. Nature 507, 62–67. https://doi.org/10.1038/nature13011

Sundaresan, R., Parameshwaran, H.P., Yogesha, S.D., Keilbarth, M.W., Rajan, R., 2017. RNA-Independent DNA Cleavage Activities of Cas9 and Cas12a. Cell Reports 21, 3728–3739. https://doi.org/10.1016/j.celrep.2017.11.100

Tao, P., Wu, X., Rao, V., 2018. Unexpected evolutionary benefit to phages imparted by bacterial CRISPR-Cas9. Science Advances 4, eaar4134. https://doi.org/10.1126/sciadv.aar4134

Tsai, S.Q., Nguyen, N.T., Malagon-Lopez, J., Topkar, V.V., Aryee, M.J., Joung, J.K., 2017. CIRCLE-seq: a highly sensitive *in vitro* screen for genome-wide CRISPR–Cas9 nuclease off-targets. Nature Methods 14, 607–614. https://doi.org/10.1038/nmeth.4278

Tsai, S.Q., Zheng, Z., Nguyen, N.T., Liebers, M., Topkar, V.V., Thapar, V., Wyvekens, N., Khayter, C., Iafrate, A.J., Le, L.P., Aryee, M.J., Joung, J.K., 2015. GUIDE-seq enables genome-wide profiling of off-target cleavage by CRISPR-Cas nucleases. Nature Biotechnology 33, 187–197. https://doi.org/10.1038/nbt.3117

Vakulskas, C.A., Behlke, M.A., 2019. Evaluation and Reduction of CRISPR Off-Target Cleavage Events. Nucleic Acid Therapeutics 29, 167–174. https://doi.org/10.1089/nat.2019.0790

Vakulskas, C.A., Dever, D.P., Rettig, G.R., Turk, R., Jacobi, A.M., Collingwood, M.A., Bode, N.M., McNeill, M.S., Yan, S., Camarena, J., Lee, C.M., Park, S.H., Wiebking, V., Bak, R.O., Gomez-Ospina, N., Pavel-Dinu, M., Sun, W., Bao, G., Porteus, M.H., Behlke, M.A., 2018. A high-fidelity Cas9 mutant delivered as a ribonucleoprotein complex enables efficient gene editing in human hematopoietic stem and progenitor cells. Nature Medicine 24, 1216–1224. https://doi.org/10.1038/s41591-018-0137-0

Vriend, L.E.M., Krawczyk, P.M., 2017. Nick-initiated homologous recombination: Protecting the genome, one strand at a time. DNA Repair 50, 1–13. https://doi.org/10.1016/j.dnarep.2016.12.005

Yan, W.X., Hunnewell, P., Alfonse, L.E., Carte, J.M., Keston-Smith, E., Sothiselvam, S., Garrity, A.J., Chong, S., Makarova, K.S., Koonin, E.V., Cheng, D.R., Scott, D.A., 2019. Functionally diverse type V CRISPR-Cas systems. Science 363, 88–91. https://doi.org/10.1126/science.aav7271

Yarrington, R.M., Verma, S., Schwartz, S., Trautman, J.K., Carroll, D., 2018. Nucleosomes inhibit target cleavage by CRISPR-Cas9 in vivo. Proc Natl Acad Sci USA 115, 9351–9358. https://doi.org/10.1073/pnas.1810062115

Zhang, L., Rube, H.T., Vakulskas, C.A., Behlke, M.A., Bussemaker, H.J., Pufall, M.A., 2020. Systematic in vitro profiling of off-target affinity, cleavage and efficiency for CRISPR enzymes. Nucleic Acids Res. https://doi.org/10.1093/nar/gkaa231

